# Natural selection and neutral mutations through the lens of viruses

**DOI:** 10.1101/2022.10.10.511542

**Authors:** Ji-Ming Chen, Guo-Hui Li, Huan-Yu Gong, Ming-Hui Sun, Yu-Fei Ji, Rui-Xu Chen, Su-Mei Tan, Ji-Wang Chen

**Affiliations:** School of Life Science and Engineering, Foshan University, Guangdong, China; Department of Medicine, University of Illinois at Chicago, Illinois, USA

## Abstract

The Neutral Theory and the Modern Synthesis, a modified version of Darwin’s theory, have been arguing for decades about the influence of natural selection on molecular evolution^1-10^. Here we elucidate through the lens of viruses that a frequently used method^11-17^ employing the ratio of nonsynonymous versus synonymous substitution rates has dramatically underestimated the influence of natural selection on molecular evolution. We also find novel evidence from viral sequences to support the co-existence of the crucial role of natural selection in molecular evolution and the ubiquity of neutral mutations. The co-existence has perplexed biologists for decades^2,5,7^. We then elucidate for the first time the causality between natural selection and the ubiquity of neutral mutations with a novel interpretation of natural selection. This novel interpretation incorporates biochemistry, genetics, epigenetics, physiology, and dynamics. It holds that natural selection acts directly on the overall phenotypic performance of organisms and indirectly on each genomic site or phenotypic trait. It highlights not only restrictions and competitions but also freedom and diversity, besides the overall harmonious development of organisms and human societies. Therefore, this novel interpretation could have far-reaching implications in the natural and social sciences.

---

Molecular evolution constitutes the genetic basis of biological evolution, which is fundamental to the natural and social sciences. Molecular evolution is affected by random genetic drift and natural selection^2,17,18^. Random genetic drift fixes selectively neutral or nearly neutral mutations by chance, while natural selection promotes beneficial mutations and inhibits deleterious mutations^19-21^.

As a modified version of Darwin′s theory, the Modern Synthesis holds that molecular evolution mainly depends on natural selection. By contrast, the Neutral Theory, which was established after the Modern Synthesis, holds that molecular evolution mainly depends on random genetic drift of neutral mutations^7-10^. Biologists have been debating these two contradictory views for decades^1-6^.

Nucleotide substitution in protein-coding genes is one of the major research topics in molecular evolution and occurs more frequently than other types of genomic changes (e.g., nucleotide insertion or deletion, gene recombination, or gene addition or deletion). Nucleotide substitution in protein-coding genes is also more straightforward than other types of genomic changes. Therefore, we employed it here to elucidate the role of natural selection in molecular evolution and the relationship between natural selection and neutral mutations.

The ratio (ω) of nonsynonymous versus synonymous substitution rates (dN/dS) has been frequently employed for over 20 years to examine the role of natural selection in nucleotide substitution at numerous sites in protein-coding genes^11-17^. The ω value of a site significantly less than 1, greater than 1, or approximately 1 respectively represents a site under negative, positive, or neutral selection, which is abbreviated as SNgS, SPS, or SNtS below. Mutations are inhibited, allowed, and promoted by negative, neutral, and positive selection, respectively.

Because many viruses are haploid with relatively simple structures and functions, and they evolve much more rapidly than cellular organisms, they are highly valuable to validate evolutionary theories or views^17,22-24^. As such we employed viral sequences to elucidate three mechanisms that the ω approach mentioned above has dramatically underestimated the influence of natural selection on molecular evolution. We also employed viral sequences to clarify the roles of natural selection and neutral mutations in molecular evolution, which has perplexed biologists for decades^2,5,7,8,25^. To simplify the relevant elucidation, neutral mutations and nearly neutral mutations are unified as neutral mutations in this report.

## The first mechanism for underestimating selection

We randomly selected 10 viral genes and randomly retrieved 500 qualified open reading frame (ORF) sequences from GenBank for each of these genes. We calculated SPSs and SNgSs in each gene using the 500 sequences, and calculated them again using 200, 200, 50, 50, 10, and 10 sequences, respectively. These sets of 200, 50, or 10 sequences were randomly selected from the 500 sequences through equidistant sampling to further reduce the influence of phylogenetic branches on the ω values. The two sets of 200 and 50 sequences did not overlap each other.

We found that the SPSs in all these 10 viral genes calculated with the Mixed-Evolutionary-Mixed-Effects (MEME) method^24^ increased with the sequences involved in the calculations (Fig. 1, Table S1), with statistical significance (*P*<0.01, by the Mann-Kendal test). For example, the percentages of SPSs in the 10 genes calculated with 50 sequences reduced by 60.8% ± 16.0% (37.9−86.7%), compared with the percentages calculated with 500 sequences.

**Fig. 1.**
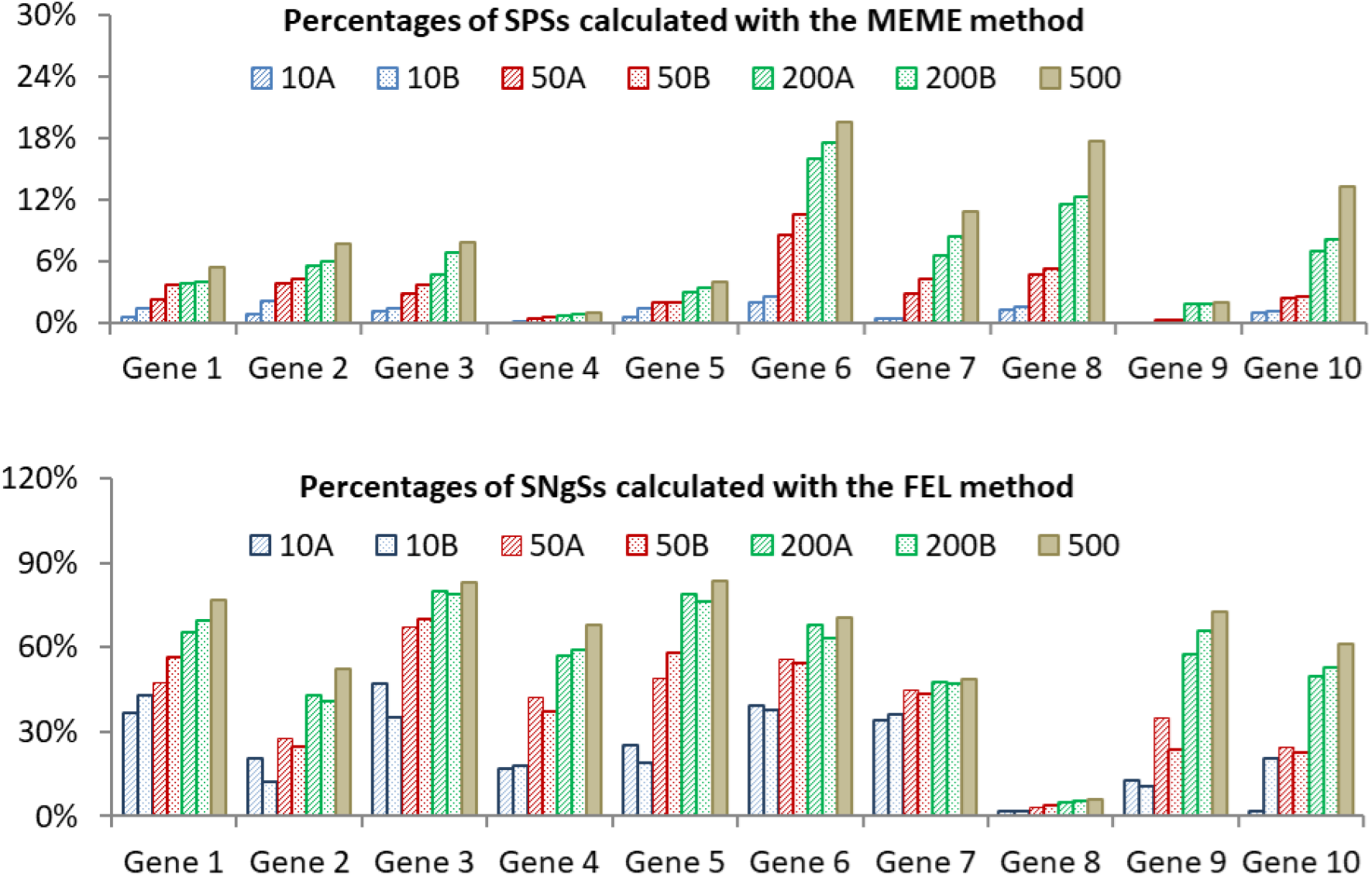
Percentages of the sites under positive selection (SPSs) and the sites under negative selection (SNgSs) in 10 viral genes calculated with 10 (10A, 10B), 50 (50A, 50B), 200 (200A, 200B) or 500 sequences.

We also found that the SNgSs in all the 10 viral genes calculated with the Fixed-Effects-Likelihood (FEL) method^24^ increased with the sequences involved in the calculations (Fig. 1, Table S1), with statistical significance (*P*<0.01, by the Mann-Kendal test). For example, the percentages of SNgSs in the 10 genes calculated with 50 sequences reduced by 37.0% ± 17.3% (7.7−67.2%), compared with the percentages calculated with 500 sequences.

We randomly checked 12 reports published in 2022 which employed 9−538 (median = 43 < 50) sequences for estimating the selection pressure on sites in protein-encoding genes of viruses using the ω approach^26-37^. Therefore, as per the above data calculated with 50 sequences using the MEME and FEL methods, at least half of these 12 reports could underestimate SPSs and SNgSs by more than 60.8% and 37.0%, respectively, in the relevant viral genes. This underestimate results from the statistical error of inadequate sampling because too few sequences can exhibit too few nucleotide substitutions for estimating the selection impact.

## The second mechanism for underestimating selection

Of the 500 qualified ORF sequences of Gene 9 (the PB2 gene of zoonotic H7N9 subtype avian influenza viruses identified after 2013 in China), 411 were from avian samples, and 89 from infected human samples). Most of the related humans were infected with the viruses through the avian-to-human transmission, which means that the viruses from human samples likely had not been fully adapted to human bodies through human-to-human transmission^38^.

\We found that the acidic amino acid residue 627E in the PB2 gene existed in 98.5% of the viruses from avian samples, and that alkaline amino acid residue 627K existed in 58.4% of the viruses from human samples (Table 1). The change suggests that the mutation E627K was beneficial and under positive selection in human bodies, but deleterious and under negative selection in avian bodies, which has been supported by various studies ^39,40^. The polymorphism of the triplet codons at site 627 shown in Table 1 suggests that this site was also under neutral selection and negative selection in human bodies: the neutral selection allowed the synonymous mutations of GAG↔GAA and AAG↔AAA; the negative selection inhibited the occurrence of the potential one-nucleotide substitutions at this site: GAG/GAA→GAT/GAC (E627D), GAG/GAA→GGG/GGA (E627G), and GAG/GAA→TAG/TAA (E627X) in both avian and human bodies (*P*<0.01, by the Chi-square test).

**Table 1.** Distribution of the triplet codons at site 627 in the PB2 gene of 500 H7N9 subtype influenza viruses isolated in China

|  | Viruses from avian samples (*n*=411) |  |  |  |  | Viruses from human samples (*n*=89) |  |  |  |  |
| --- | --- | --- | --- | --- | --- | --- | --- | --- | --- | --- |
| Residue | E | E | K | K | V | E | E | K | K | V |
| Codon | GAG | GAA | AAG | AAA | GTG | GAG | GAA | AAG | AAA | GTG |
| Count | 275 | 130 | 4 | 1 | 1 | 26 | 10 | 44 | 8 | 1 |
| Rate (%) | 66.9 | 31.6 | 1.0 | 0.2 | 0.2 | 29.2 | 11.2 | 49.4 | 9.0 | 1.1 |

The above data show that site 627 was under positive, neutral, and negative selection simultaneously in human bodies, and under neutral and negative selection simultaneously in avian bodies. This example suggests the second mechanism that the ω approach has underestimated nature selection in molecular evolution: The ω approach assumes each genomic site is under only one type of selection, neglects the possibility that a genomic site can be under two or three types of selection in the same branch.

The MEME, FEL, Unconstrained-Bayesian-Approximation (FUBAR), and Single-Likelihood-Ancestor-Counting (SLAC) methods in the ω approach all assume that nonsynonymous substitutions of a site are equally distributed among possible nonsynonymous substitutions^11-17,24^. This assumption is incorrect for some genomic sites from statistics and from the biological perspective, as supported by the fact that numerous genomic sites favored only certain amino acid residues (e.g., the above example of site 627 in the PB2 gene likely favored only E in avian bodies and only K in human bodies)^39,40^. These sites were termed sites under directional selection, and they are under positive, neutral, and negative selection simultaneously^22,23^. This further clarifies the second mechanism for underestimating natural selection in molecular evolution.

## The third mechanism for underestimating selection

The third mechanism that the ω approach has underestimated positive selection in molecular evolution has been reported^24^. The ω approach sometimes neglects the sites under positive selection in some phylogenetic branches and under negative selection in some other phylogenetic branches. Therefore, these sites under natural selection can be calculated as SNgSs or SNtSs if their ω values are calculated with a pervasive method, such as FEL, FUBAR, and SLAC^24^.

This type of underestimate can be partially circumvented using the MEME method which can identify SPSs from an episodic view^24^. Table 2 showed approximately 69.7% ± 28.8% (20.3−100.0%), 76.7% ± 27.0% (25.3−100.0%), and 71.7% ± 25.3% (25.6−100.0%) of the SPSs in the 10 viral genes calculated with the MEME were not calculated as SPSs by the FEL, FUBAR, and SLAC methods, respectively. Moreover, the SPSs calculated with the pervasive methods could be unstable if using different sets of data, as shown by the examples given in Table 3 where, for each gene, the SPSs calculated with two sets of non-overlapping 200 sequences randomly selected from 500 sequences overlapped approximately 25.6% ± 26.9% (0.0−81.5%). Because the two sets of non-overlapping 200 sequences were randomly selected, the differences shown in Table 3 did not result from different selection among phylogenetic branches.

**Table 2.** Counts of sites under positive selection (SPSs) calculated with four methods using 500 sequences per gene.

| Method | Gene |  |  |  |  |  |  |  |  |  |
| --- | --- | --- | --- | --- | --- | --- | --- | --- | --- | --- |
|  | 1 | 2 | 3 | 4 | 5 | 6 | 7 | 8 | 9 | 10 |
| MEME | 31 | 18 | 44 | 10 | 20 | 39 | 23 | 71 | 15 | 179 |
| FEL | 7 | 5 | 6 | 6 | 0 | 30 | 2 | 63 | 1 | 13 |
| FUBAR | 4 | 3 | 4 | 3 | 0 | 27 | 3 | 59 | 1 | 2 |
| SLAC | 9 | 6 | 8 | 4 | 0 | 29 | 2 | 51 | 1 | 14 |

**Table 3.**
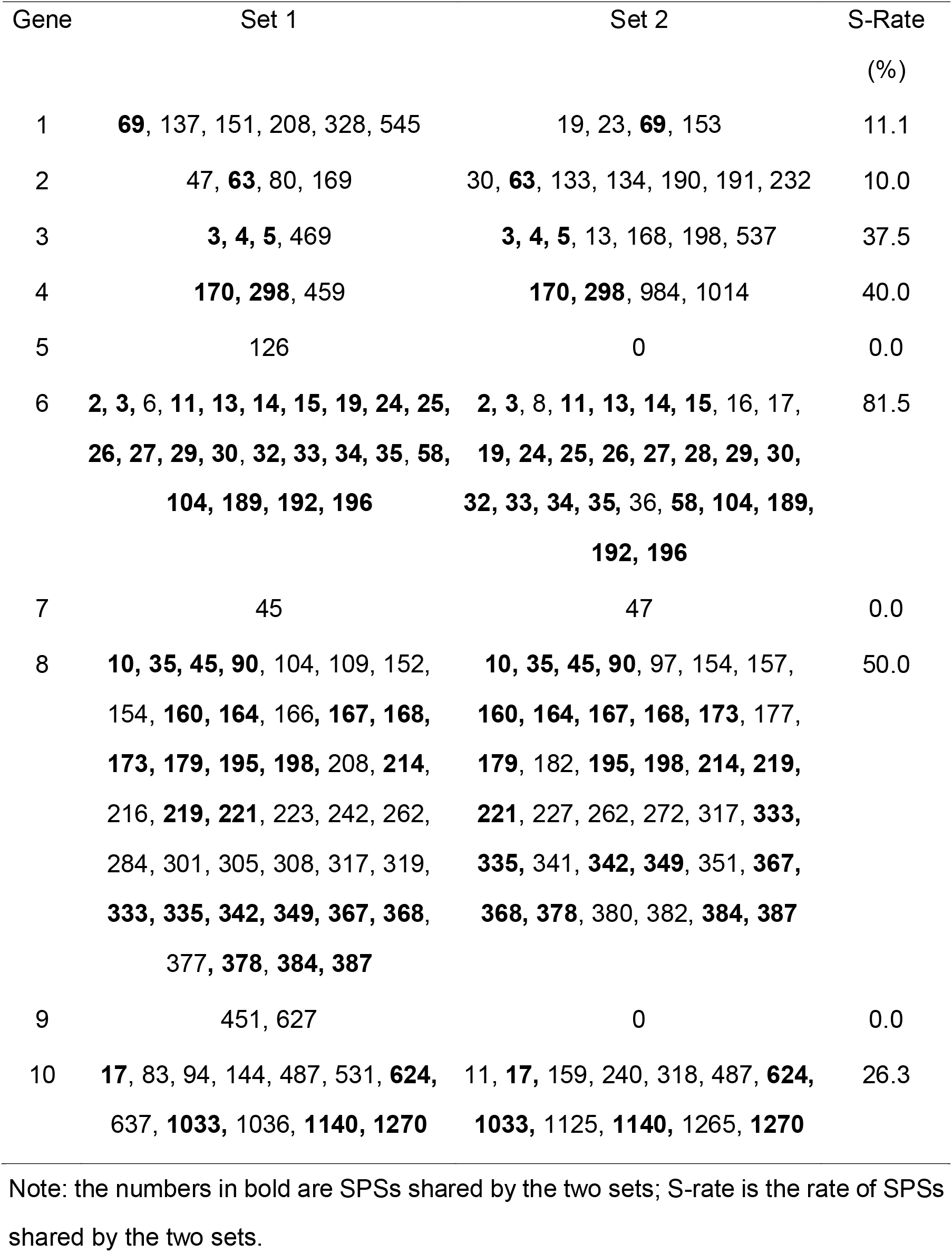
The Sites under positive selection (SPSs) calculated with the FEL method and two non-overlapping sets of 200 sequences per gene.

## Ubiquity of natural selection and neutral mutations

Fig. 1 and Table S1 suggest that approximately 69.6% ± 16.5% (35.8−91.5%) and 8.9% ± 6.4% (0.9−19.5%) of all the sites in the 10 viral genes were under negative and positive selection, respectively, if calculated using 500 sequences for each gene with the MEME method to avoid the above first and third mechanisms for underestimating selection. Additionally, approximately 3.6% ± 2.8% (0.5−8.0%) of all the sites in the genes encountered no nucleotide substitutions among their 500 sequences, and these sites could be under negative selection that inhibited both synonymous and nonsynonymous mutations. These data demonstrate that natural selection is crucial to the evolution of these 10 viral genes, as approximately 82.1% (69.6% + 8.9% + 3.6%) of all the sites in the genes were under negative or positive selection. When the second mechanism underestimating selection mentioned above is considered, more than 82.1% sites in the genes could be under negative or positive selection. This is consistent with many studies that support the crucial role of nature selection in molecular evolution^2,3^.

Approximately 66.6% ± 14.3% (46.8−82.7%) of all the nucleotide substitutions in the 10 viral genes we studied were synonymous mutations, as identified with the SLAC method (Fig. 2). Meanwhile, synonymous mutations occurred at least at SNtSs and SNgSs, which totally covered approximately 92.0% ± 7.3% (77.8−98.6%) of all the sites in the viral genes (Table S1). Therefore, consistent with previous studies^2,3^, these data demonstrate that neutral mutations are ubiquitous in the 10 viral genes because synonymous mutations fixed in the genomes are usually selectively neutral, although some synonymous mutations are deleterious and cannot be fixed due to negative selection^41^. Because some non-synonymous mutations fixed in the genomes, particularly for those at SNtSs, could be also selectively neutral, >66.6% nucleotide substitutions in the 10 viral genes were likely neutral in selection.

**Fig. 2.**
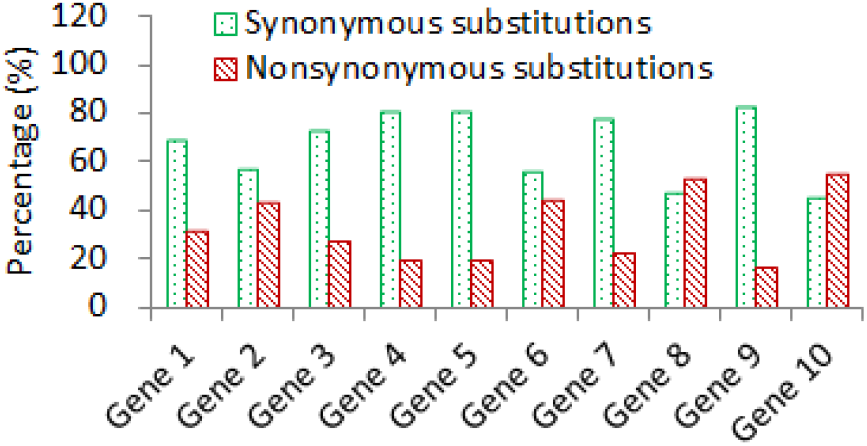
Percentages (%) of the synonymous and nonsynonymous substitutions in 10 viral genes calculated with the SLAC method and 500 sequences per gene.

## Novel interpretation of natural selection

To explain the co-existence of the ubiquity of neutral mutations and natural selection in molecular evolution, we provide a novel interpretation of natural selection which incorporates well-known knowledge in biochemistry, genetics, epigenetics, physiology, and dynamics (Fig. 3).

**Fig. 3.**
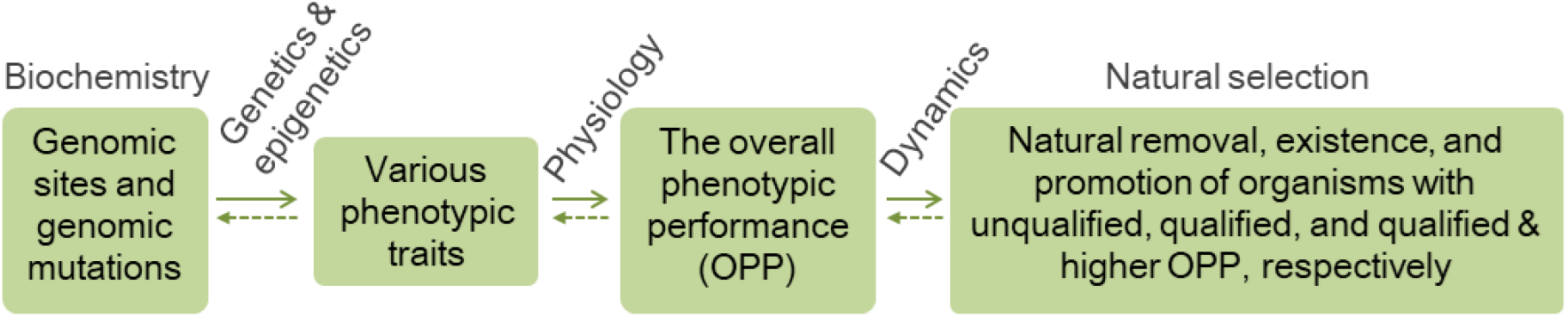
The formation (→) and action (←) of natural selection.

In biochemistry, minor or major variations at any genomic sites are unavoidable, which leads to genetic diversity, and each genomic site can have a few statuses which largely correspond to mutations or alleles at the population level.

In genetics, due to a plenty of physical and chemical reactions under the direction of the information encoded in the genomic sequences, an organism obtains numerous phenotypic traits. Meanwhile, minor genomic mutations can cause minor to major heritable changes in phenotypic traits, so can major genomic mutations^5^. Epigenetic changes can also lead to changes in phenotypic traits.

In physiology, numerous phenotypic traits collectively constitute the overall phenotypic performance (OPP) associated with the survival and reproduction of the organism. In mammals, the OPP includes the ability to obtain food, recover from infections, defeat enemies, and win mating opportunities; in viruses, the OPP includes the ability to survive in the environment, infect hosts, replicate in hosts, and spread among hosts. Many phenotypic traits, genomic mutations, and epigenetic changes can individually disqualify and cannot qualify the OPP (e.g., a genomic mutation associated with the heart can disqualify and cannot qualify the OPP).

In dynamics, only those organisms with qualified OPP can survive and reproduce themselves, and those organisms with qualified and better OPP can survive and reproduce themselves more efficiently. This leads to natural selection, which means accumulation of the phenotypic traits, genomic mutations, and epigenetic changes that are beneficial to the OPP, inhibit of the phenotypic traits, genomic mutations, and epigenetic changes that are deleterious to the OPP, and random drift of the phenotypic traits, genomic mutations, and epigenetic changes that are neutral to the OPP. These dynamic changes suggest that natural selection directly acts on the OPP of organisms and indirectly on each genomic mutation, phenotypic trait, and epigenetic change.

Also in dynamics, with the most beneficial mutations accumulated in more and more genomic sites and deleterious mutations inhibited continuously at all genomic sites, more and more genomic sites can encounter almost only neutral mutations (e.g., synonymous substitutions) that can be fixed in the population. This leads to the increase in the proportion of neutral mutations and the ubiquity of neutral mutations.

For example, consistent with various studies ^39,40^, Table 1 suggests that the amino acid residue E is likely the fittest amino acid residue at site 627 in the PB2 gene of the H7N9 viruses isolated from avian samples. This means that no beneficial mutations, but neutral or deleterious mutation can occur on this site after 627E has become ubiquitous in the PB2 gene of the viruses from birds. Furthermore, hitchhiking of neutral mutations along with beneficial mutations is facilitated directly by positive selection^2,6,21^. Therefore, there is a dialectical causality between the crucial role of natural selection in molecular evolution and the ubiquity of neutral mutations.

Under natural selection, three factors promote biodiversity that includes genetic diversity and phenotypic diversity. First, as natural selection directly “selects” organisms as per their OPP, and some organisms weaker in a phenotypic trait but stronger in another phenotypic trait can have qualified OPP, some deleterious mutations (e.g., those associated with thalassemia) have been fixed in populations. Second, as a mutation neutral or deleterious to the OPP in some niches (e.g., the above E627K mutation in birds) could be beneficial to the OPP in some other niches (e.g., the above E627K mutation in humans), this mutation can be fixed in the population. Third, habitable environments promote biodiversity because it is easier for organisms to have qualified OPP in habitable environments than in harsh environments.

The above novel interpretation of natural selection is different from previous interpretations in multiple aspects (Table 4). It emphasizes selection on the OPP of organisms, encompasses both minor and major genomic, epigenetic, and phenotypic changes, underlines both competitions and development of diversity, and explains the co-existence and causality between natural selection and the ubiquity of neutral mutations, while previous interpretations emphasize selection on a single genomic or phenotypic change, reluctant to accept major genomic, epigenetic, or phenotypic changes, usually underline stringent competitions of natural selection (as mirrored in the slogan of “survival of the fittest”), and assume that natural selection and the ubiquity of neutral mutations are contradictory.

**Table 4.** Different interpretations of natural selection

| The interpretation in this report | Previous interpretations |
| --- | --- |
| Acknowledges the existence of major mutations | Neglect the existence of major mutations |
| Acknowledges the existence of major phenotypic changes | Neglect the existence of major phenotypic changes |
| Able to incorporate epigenetic changes | Unable to incorporate epigenetic changes |
| Natural selection acts directly on the overall phenotypic performance | Natural selection acts directly on a single mutation or phenotypic change |
| Natural selection leads to the ubiquity of neutral mutations | Unable to explain the ubiquity of neutral mutations |
| Able to explain genetic or phenotypic defects of organisms | Unable to explain genetic or phenotypic defects of organisms |
| Underlines competitions and diversity | Underlines competitions |
| Natural selection is stringent and lenient simultaneously | Natural selection is very stringent |

## Discussion

In this study, we first elucidate through the lens of viruses three mechanisms that dramatically underestimate natural selection in molecular evolution in the past. The first mechanism could underestimate the SPSs and SNgSs in the 10 viral genes by approximately 60.8% and 37.0% if they are calculated with 50 sequences per gene, compared with the calculation using 500 sequences per gene (Table 1). The third mechanism could underestimate the SPSs in the 10 viral genes can be underestimated by approximately 69.7% if calculated using the FEL method with 500 sequences per gene, compared with the calculation using the MEME method (Table 2). It remains unknown how much the second mechanism could underestimate natural selection in molecular evolution. Although the ω approach has underestimated the selection impact dramatically, this approach has contributed greatly to the estimate of natural selection and can be optimized to circumvent the underestimate.

We then elucidate the co-existence of the ubiquity of neutral mutations and the crucial role of natural selection in molecular evolution. We provide a novel interpretation of natural selection to explain their co-existence and causality (Fig. 3). The novel interpretation is based on well-known knowledge in biochemistry, genetics, epigenetics, physiology, and dynamics (Fig. 3). It is not solely based on the above data of the 10 viral genes. It incorporates background selection, hitchhiking selection, and simultaneous selection of various genomic mutations and phenotypic traits from a panorama view.

Natural selection is associated with restrictions and competitions, and mutations are associated with freedom and diversity. As we know, in biospheres and human societies, freedom is widely inhibited and supported simultaneously by restrictions, and diversity is widely inhibited and supported simultaneously by competitions. Furthermore, in dynamics, restrictions and competitions can lead to ubiquity of neutral freedom and neutral diversity (neutral freedom or neutral diversity represent some changes as advantageous as their original forms in the relevant restrictions or competitions). These phenomena, in turn, support the causality between the ubiquity of neutral mutations and the crucial role of natural selection in molecular evolution.

The Neutral Theory first proposed by Motoo Kimura in 1968 holds two key views: most genomic mutations fixed in the genomes are selectively neutral; natural selection is less important than random genetic drift to molecular evolution. We elucidate in this report that natural selection in molecular evolution has been dramatically underestimated, and that approximately 82.1% sites of 10 randomly selected viral genes were under negative or positive selection even with the dramatic underestimate. Therefore, we find novel evidence from viruses supporting the crucial role of natural selection in molecular evolution and negate the second view of the Neutral Theory. Meanwhile, approximately 66.6% substitutions in these genes were synonymous neutral mutations. This supports the first key view of the Neutral Theory.

Viruses cause numerous infectious diseases in humans, animals, and plants, such as COVID-19 in humans, avian influenza in animals, and rice dwarf disease in plants^42^. SPSs and SNgS in viral genes are usually important for the virus to infect its hosts, replicate in its hosts, or escape from host immunity^17^. Therefore, it is valuable to identify more authentic SPSs and SNgS in various viral genes in this report, which suggests that many more sites in the genomes of various viruses are under positive or negative selection than in previous reports.

In summary, we elucidate in this report through the lens of viruses that natural selection in molecular evolution has been dramatically underestimated. Furthermore, we show that most mutations in randomly selected viral genes are likely selectively neutral, and most sites in these genes are likely under negative or positive selection. We then elucidate the causality between natural selection and the ubiquity of neutral mutations with a novel interpretation of natural selection. This novel interpretation incorporates biochemistry, genetics, epigenetics, physiology, and dynamics. It holds that natural selection acts directly on the overall phenotypic performance of organisms. It highlights restriction and freedom, competition and diversity, as well as the overall harmonious development of organisms and human societies. This novel interpretation could have far-reaching implications in the natural and social sciences.

## Materials and Methods

### Sequence recombination detection

Sequence recombination was detected using the software tool RDP (version 4.0) with the methods of RDP, GENECONV, Chimera, MaxChiu, Bootscan, and SISCAN ^43^. The analysis was performed with default settings, and sequence recombination were judged as those detected by at least two methods with *P* < 0.05.

### Sequence selection and alignment

Nucleotide sequences encoding the whole-length open reading frame (ORF) of randomly selected 10 viral genes (Genes 1−10) were retrieved from the GenBank. Those sequences containing the ambiguous sites of N, X, M, Y, W, S, or R, or fully identical with another sequence were removed. Those sequences potentially recombined from other retrieved sequences were also removed. To minimize the influence of phylogenetic branches on the ω values, sequences of small clades distant in phylogenetics from other clades were excluded. The remaining sequences were qualified for this study and aligned using the online software tool MAFFT as per the advanced setting of G-INS-I which is slow and progressive with an accurate guide ^44^.

### Names of viral genes

These ten viral genes are HA gene of H3N2 subtype human influenza virus in *Orthomyxoviridae* (Gene 1), ORF2 gene of porcine circovirus 2 in *Circoviridae* (Gene 2), HA gene of H9N2 subtype avian influenza virus in *Orthomyxoviridae* (Gene 3), M gene of severe fever with thrombocytopenia syndrome virus in *Phenuiviridae* (Gene 4); env gene of Japanese encephalitis virus in *Flaviviridae* (Gene 5), ORF5 gene of porcine reproductive and respiratory syndrome virus 2 in *Arteriviridae* (Gene 6), polyprotein gene of Foot and mouth disease virus in *Picornaviridae* (Gene 7), S gene of hepatitis B virus in *Hepadnaviridae* (Gene 8), PB2 gene of H7N9 subtype human influenza virus in *Orthomyxoviridae* (Gene 9), and S gene of severe acute respiratory syndrome-related coronavirus in *Coronaviridae* (Gene 10).

### Calculation of selection on sites

SPSs were calculated using the online software tool Datamonkey with the method of Mixed Evolutionary Mixed Effects Model (MEME) ^24,45^. SNgSs were calculated using the same tool with the methods of Fixed Effects Likelihood (FEL), if not specified. The SPSs, SNgSs, and SNtSs in the genes were also calculated using the methods of Fast, Unconstrained Bayesian Approximation (FUBAR)^46^ and Single-Likelihood Ancestor Counting (SLAC) ^12^. Synonymous mutations and non-synonymous mutations in the genes were counted using the SLAC method. The statistical possibility (*P*) value less than 0.1 was considered to be statistically significant, except for the FUBAR method where the posterior probability greater than 0.9 was considered to be statistically significant, if not specified.

## Author contributions

J.M.C. conceived, designed, and supported this study, analyzed the data, and drafted the manuscript. J.W.C. jointly made the core conclusion, analyzed the data, and revised the manuscript. G.H.L, H.Y.G, R.X.C, S.M.H, Y.F.J, and T.S.M. collected and analyzed relevant data and revised the manuscript.

## Acknowledgements

This study was supported by the High-Level Talent Fund of Foshan University (no. 20210036). The funder did not play any role in this study.

## Data availability statement

The data supporting the views of this analysis are provided as **Tables S2** and **S3** and available from the corresponding author on request.

