## Supplementary material for "Natural selection and neutral mutations through the lens of viruses": Table S1

**Table S1**. Percentages (%) of Sites under Positive Selection (SPSs) Calculated with the MEME method, Sites under Negative Selection (SNgSs), Sites under Neutral Selection (SNtSs), and Sites without Changes (SnoCs) Calculated the FEL method.

| Sequences employed | Gene 1 | | | | Gene 2 | | | | Gene 3 | | | | Gene 4 | | | | Gene 5 | | | |
|---|---|---|---|---|---|---|---|---|---|---|---|---|---|---|---|---|---|---|---|---|
| | SPS | SNgS | SnoC | SNtS | SPS | SNgS | SnoC | SNtS | SPS | SNgS | SnoC | SNtS | SPS | SNgS | SnoC | SNtS | SPS | SNgS | SnoC | SNtS |
| 10 | 0.5 | 36.7 | 31.1 | 31.8 | 0.9 | 12.3 | 66.8 | 20.9 | 1.1 | 47.3 | 21.8 | 30.7 | 0.0 | 17.0 | 65.0 | 18.1 | 0.6 | 25.4 | 43.2 | 31.4 |
| 10 | 1.4 | 42.9 | 27.6 | 29.2 | 2.1 | 20.4 | 61.3 | 17.9 | 1.4 | 35.2 | 28.9 | 35.5 | 0.2 | 18.2 | 62.0 | 19.9 | 1.4 | 18.8 | 50.4 | 30.8 |
| 50 | 2.3 | 47.5 | 15 | 36.6 | 3.8 | 27.7 | 44.7 | 26.8 | 2.9 | 67.5 | 8 | 24.1 | 0.4 | 42.5 | 31.7 | 25.7 | 2.0 | 49.2 | 21.0 | 29.8 |
| 50 | 3.7 | 56.7 | 11.5 | 31.4 | 4.3 | 24.7 | 42.1 | 31.9 | 3.8 | 69.8 | 6.6 | 22.9 | 0.6 | 37.0 | 36.5 | 26.4 | 2.0 | 58.0 | 12.8 | 29.0 |
| 200 | 3.9 | 65.5 | 5.8 | 27.6 | 5.5 | 43 | 17.4 | 37.9 | 4.6 | 80.2 | 2.9 | 16.3 | 0.7 | 57.0 | 15.5 | 27.2 | 3.0 | 79.0 | 2.2 | 18.6 |
| 200 | 4.1 | 69.8 | 4.8 | 24.7 | 6 | 40.9 | 15.7 | 40.4 | 6.8 | 78.9 | 3.2 | 16.6 | 0.8 | 59.3 | 14.8 | 25.5 | 3.4 | 76.2 | 4.2 | 19.6 |
| 500 | 5.5 | 77 | 3.4 | 18.4 | 7.7 | 52.3 | 4.3 | 41.3 | 7.9 | 83.2 | 1.1 | 14.6 | 0.9 | 67.8 | 7.5 | 24.0 | 4.0 | 83.8 | 1.2 | 15.0 |

| Sequences employed | Gene 6 | | | | Gene 7 | | | | Gene 8 | | | | Gene 9 | | | | Gene 10 | | | |
|---|---|---|---|---|---|---|---|---|---|---|---|---|---|---|---|---|---|---|---|---|
| | SPS | SNgS | SnoC | SNtS | SPS | SNgS | SnoC | SNtS | SPS | SNgS | SnoC | SNtS | SPS | SNgS | SnoC | SNtS | SPS | SNgS | SnoC | SNtS |
| 10 | 2 | 39.5 | 23 | 35.5 | 0.5 | 64.3 | 13.6 | 21.6 | 1.3 | 10.3 | 60.3 | 28.8 | 0 | 12.5 | 70 | 17.5 | 1 | 1.9 | 5.8 | 91.8 |
| 10 | 2.5 | 38 | 25 | 33.5 | 0.5 | 67.6 | 12.2 | 19.7 | 1.5 | 11.8 | 52.8 | 35.3 | 0 | 10.7 | 68 | 21.3 | 1.2 | 20.3 | 5.8 | 73.6 |
| 50 | 8.5 | 56 | 8.5 | 29 | 2.8 | 84.5 | 3.8 | 11.7 | 4.8 | 20.8 | 35.5 | 39.8 | 0.3 | 35.2 | 34.4 | 30.3 | 2.4 | 24.7 | 5.8 | 69.3 |
| 50 | 10.5 | 54.5 | 9 | 28 | 4.2 | 81.2 | 1.9 | 16.9 | 5.3 | 21.8 | 31.0 | 43.0 | 0.3 | 23.8 | 41.2 | 34.9 | 2.5 | 22.6 | 5.8 | 71.2 |
| 200 | 16 | 68 | 3.5 | 16.5 | 6.6 | 89.7 | 0.9 | 8.9 | 11.5 | 27.8 | 18.8 | 43.3 | 1.8 | 57.7 | 14.8 | 27.3 | 6.9 | 49.5 | 5.6 | 44 |
| 200 | 17.5 | 63.5 | 5 | 18 | 8.5 | 88.7 | 0.9 | 9.9 | 12.3 | 31.8 | 14.5 | 44.5 | 1.8 | 66.1 | 12.9 | 20.9 | 8.2 | 52.8 | 5.8 | 40.4 |
| 500 | 19.5 | 70.5 | 3 | 11.5 | 10.8 | 91.5 | 0.5 | 7 | 17.8 | 35.8 | 8.0 | 40.5 | 2 | 72.6 | 7.9 | 19.4 | 13.2 | 61 | 5.6 | 32.4 |
